# Risk and Loss Aversion and Attitude to COVID and Vaccines in Anxious Individuals

**DOI:** 10.1101/2023.05.05.539520

**Authors:** Filippo Ferrari, Jesse Alexander, Peggy Seriès

**Affiliations:** Institute for Adaptive and Neural Computation School of Informatics, University of Edinburgh, United Kingdom

**Keywords:** Anxiety, Risk aversion, Loss aversion, COVID-19, Computational Psychiatry

## Abstract

Anxious individuals are known to show impaired decision-making in economic gambling task and in everyday life decisions. This impairment can be due to aversion to uncertainty about outcomes (risk aversion) and/or aversion to negative outcomes (loss aversion). We investigate how non-clinical individuals with high levels of Generalised Anxiety Disorder (GAD) (N = 54) behave compared to less anxious subjects (N = 61) in a gambling decision-making task delivered online and designed to separate the distinct influences of risk and loss aversion on decision-making. By modelling subjects’ choices using computational models derived from Prospect Theory and fitted using Hierarchical Bayesian methods, we estimate individual levels of risk and loss aversion. Moreover we analyse the reaction times and choice data using Hierarchical Drift Diffusion Models (HDDMs). We also link estimates of these parameters to individual propensity to risk averse behaviours during the COVID pandemic, like wearing safer types of face masks, or completing a COVID vaccination course. We report increased loss aversion in individuals with increased level of GAD compared to less anxious individuals and no differences in risk aversion. The HDDM analysis suggests that high anxiety subjects have a reduced bias towards gambling in trials where there is a risk to loose points, also consistent with them being loss averse. We also report no links between risk and loss aversion and attitudes towards COVID and vaccines. These results shed new light on the interplay of anxiety and risk and loss aversion and they can provide useful directions for clinical intervention.

## 1 Introduction

Anxiety disorders have long been associated with behavioural differences in decision-making (Bishop and Gagne, 2018). Here we examine economic decisions in which two main factors are at play: risk and loss aversion. A famous behavioural economics theory called Prospect Theory (Kahneman and Tversky, 1979) describes how humans prefer sure outcomes with lower magnitude (low risk) compared to less sure outcomes with higher magnitude (high risk), which results in risk averse behaviours. Prospect Theory also describes how individuals are generally considered loss averse, i.e. humans prefer to avoid losses to acquiring gains of equivalent value, usually with a 2:1 ration between losses and gains.

In this study we use computational models derived from Prospect Theory to quantify the influence of risk and loss aversion on human decision-making. We adapt a gambling task previously used by Charpentier et al. (2017) to be administered online on a non-clinical sample of adults. This task is particularly interesting as it has been one the first of its kind which allows to separate the influence of risk aversion (i.e., aversion to uncertainty about outcomes) and loss aversion (i.e., aversion to negative outcomes) on economic decision-making.

The impact of anxiety on risk aversion has been studied in several studies generally linking increased level of anxiety to increased risk aversion biases in decision-making tasks (Maner et al., 2007; Mueller et al., 2010; Giorgetta et al., 2012; Charpentier et al., 2017; see Hartley and Phelps, 2012; Phelps, Lempert, and Sokol-Hessner, 2014; Bishop and Gagne, 2018; Wake, Wormwood, and Satpute, 2020, for reviews). Despite this, results from studies employing computational models are mixed. As discussed above, Charpentier et al. (2017) reports increased risk aversion in anxious individuals, whereas Xu et al. (2020) reports no difference in risk aversion for low and high anxiety groups. Two recent studies (Wise et al., 2022; Pike et al., 2023) also failed to observe a relationship between anxiety and differences in risk-taking, despite using different decision-making tasks. A recent meta-analysis (Wake, Wormwood, and Satpute, 2020) on the effects of fear (and anxiety) on risk-taking reports a small to moderate effect size linking fear to increased risk aversion, but with high heterogeneity in effect sizes.

Loss aversion, on the other hand, has only been investigated in anxiety in a handful of studies. Charpentier et al. (2017) found that individuals with high anxiety did not show differences in loss aversion compared to the low anxiety individuals, consistent with some previous work in the literature (for a review: (Sediyama, de Castro Martins, and Martins Teodoro, 2020)). Another study by Xu et al. (2020) reports increased loss aversion and no differences in risk aversion in high anxious individuals compared to low anxious subjects.

Both risk and loss aversion are important factors in anxiety with major effects, not only on obvious economic decisions, but also on individuals’ decisions in everyday life. This is particularly true in crises situation such as observed during the COVID pandemic. When the health of large populations is at stake, it is important to understand what motivates specific attitudes and behaviours toward vaccinations, preemptive measures like wearing face masks and adherence to government guidelines. For example, intention to be vaccinated was associated with more positive beliefs around COVID vaccines and reduced beliefs that these would lead to negative side effects or be unsafe (Sherman et al., 2021). A recent longitudinal study (Shou et al., 2022) showed that risk averse individuals in Australia were more likely to act in a way that would reduce their risk of contracting COVID-19. At the same time, anxious individuals showed increased avoidance in implementing changes in their actions as the pandemic developed, for example continuing to cancel family events or trips even as the pandemic eased off. The first result is also consistent with Wise et al. (2022), where it is reported that higher risk aversion is linked to higher perceived threat from the pandemic. In our study, we are interested in determining whether behaviours like receiving COVID vaccinations, following government’s guidelines, wearing more protective types of face masks are modulated by anxiety and individual propensity to taking risks and/or accepting losses. We hypothesise that individual who adopted safer attitudes and behaviours around COVID would show increased risk aversion, and that this difference is modulated by anxiety.

Here, we also model choice and reaction data using Hierarchical Drift Diffusion Models (HDDMs) (Wiecki, Sofer, and Frank, 2013), a hierarchical Bayesian extension to a type of sequential sampling model used to model quick binary decisions. The standard model used in this study (Ratcliff, 1978) assumes that, at each trial, evidence is accumulated until it reaches on of two decision thresholds which in turn triggers a response. This model allows to discriminate between several decision components: the boundary separation between the two options, the starting point between the boundaries representing decision biases, the drift rate which represents the rate of evidence accumulation and the non-decision time which represents the time at the start of the trial in which the agent is processing other information (for example stimuli encoding). It is generally believed that high levels of anxiety result in faster evidence accumulation for ‘threatening’ choices in studies employing Drift Diffusion Models (DDMs) (see Roberts and Hutcherson, 2019, for a review). Few studies investigated risk and/or loss aversion using DDMs, but one study (Clay et al., 2017) conducted on healthy subjects using a cards-based gambling task found that increased levels of loss aversion are linked to reduced drift rates when subjects are presented with a deck of cards with higher probability of gaining points, rather than a deck with higher probability of losing them, indicating that high loss averse individuals accumulate evidence more slowly, even when the possibility of winning is higher.

In summary, this work has therefore several goals. First, we replicate and extend, using an online version of the same gambling task, previous work (Charpentier et al., 2017) linking anxiety to increased risk aversion. Second, we investigate how individual levels of anxiety as well as risk and loss aversion relate to decision-making in the context of the COVID pandemic.

## 2 Methods

### 2.1 Ethics

The study was approved by the Informatics Research Ethics Process (application number 2019/84385) of the School of Informatics, University of Edinburgh. All participants gave informed consent.

### 2.2 Participants

We recruited 163 participants based in the United Kingdom (80 with no COVID vaccinations, 83 with 2 or more doses of an approve COVID vaccine) in the months of September and October 2022 through Prolific. Participants’ data was anonymised by Prolific and further anonymised after data collection. Participants were selected from the Prolific users residing in the United Kingdom and who received either 0 or 1+ dose of an approved COVID vaccine. Participants were paid £2.67 (the task took on average 20 minutes to complete, corresponding to a pay of £8/hr) with the possibility to win up to £1 proportional to their performances during the task. We performed a power analysis based on the results reported by Charpentier et al. (2017): a sample of 104 participant is required for risk aversion being higher in anxious individuals relative to control subjects in order to achieve an effect size of Cohen’s d = 0.72 at a significance level of 0.05 and with power of 0.95. We employed exclusion criteria based on attention checks and on the participants behaviour, these are described in detail in the Supplementary Information.

After applying these exclusion criteria, we conducted the experiments on 115 subjects, for whom we report demographic information in table 1, together with the questionnaire results described in subsection 2.4.

**Table 1:** Sample demographics and questionnaire results. Values are mean (SD). The last row refers to COVID Score group differences between the vaccinated and unvaccinated groups when the number of vaccines is not included in the score.

| | All | Vaccinated | Unvaccinated | $t_{113}$ | $p$-value |
| --- | --- | --- | --- | --- | --- |
| Women:Men (Sum) | 59:56 | 37:23 (60) | 22:33 (55) | - | - |
| Age in years | 39.69 (12.77) | 40.88 (13.51) | 38.38 (11.90) | 1.049 | 0.296 |
| STAI State Anxiety | 40.87 (13.30) | 43.01 (13.83) | 38.54 (12.38) | 1.819 | 0.071 |
| STAI Trait Anxiety | 48.18 (13.87) | 47.86 (14.04) | 48.52 (13.79) | -0.254 | 0.799 |
| GAD-7 | 7.28 (5.73) | 7.83 (5.71) | 6.67 (5.74) | 1.085 | 0.279 |
| COVID Score | 41.12 (13.79) | 50.88 (8.62) | 30.47 (9.94) | 11.787 | < **0.001** |
| COVID Score (No #vax) | 38.65 (12.71) | 47.06 (8.42) | 29.47 (9.94) | 10.267 | < **0.001** |

Throughout this work we refer as the low/high anxiety group as the lower/higher median split of the data using GAD-7 scores (median GAD-7: 7) and as the vaccinated group as subjects who received 1 or more doses of a COVID-19 approved vaccine (high anxiety and vaccinated: *N* = 30; low anxiety and vaccinated: *N* = 30; high anxiety and unvaccinated: *N* = 24; low anxiety and unvaccinated: *N* = 31).

### 2.3 Task

The task was adapted from Charpentier et al. (2017) and implemented using jsPsych (de Leeuw, 2015). It could only be completed using a computer. During the gambling task participants are competing in a game show where, at each trial, they are asked to make a choice between a gambling wheel and a sure option. Each option on the gambling wheel has a 50% probability to occur. Example trials are shown in fig. 1. There are two types of trials that are shown to the participants, *mixed gamble* trials and *gain-only* trials. In mixed gamble trials one option leads to a sure option of £0, whereas the other option consists of a wheel with a 50% chance to win a monetary reward, and a 50% chance to lose some of the money accumulated so far. In gain-only trials the sure option consists of a fixed, small amount of money, whereas the other option consist of a wheel with a 50% chance to win a larger monetary reward, and a 50% chance to win nothing. During gain-only trials, only risk aversion contributes to choosing from the safe option compared to the gambling option, while both risk and loss aversion contribute to choosing from the safe option in mixed gamble trials. At the beginning of the experiment, participants are instructed to win as much money as possible, and they are told that they will receive a bonus payment through Prolific based on their performance during the experiment. A practice section consisting of 40 trials is used to introduce the participants to the task and to adapt, through a double staircase procedure, the amounts shown at each trial to a range of expected values centered on the subject’s indifference point, i.e. the expected value corresponding to an accept rate of 50% between the gamble and the sure option. The main section of the experiment consists of 148 trials (98 mixed gamble trials and 50 gain-only trials in random order, the same proportion of mixed gamble/gain-only used in Charpentier et al.), in which participants have to select an option within 4s, followed by a 2s animation showing the sure/win/loss amount. Participants were instructed to use keys [A] and [L] of their keyboards to choose between options.

**Figure 1:**
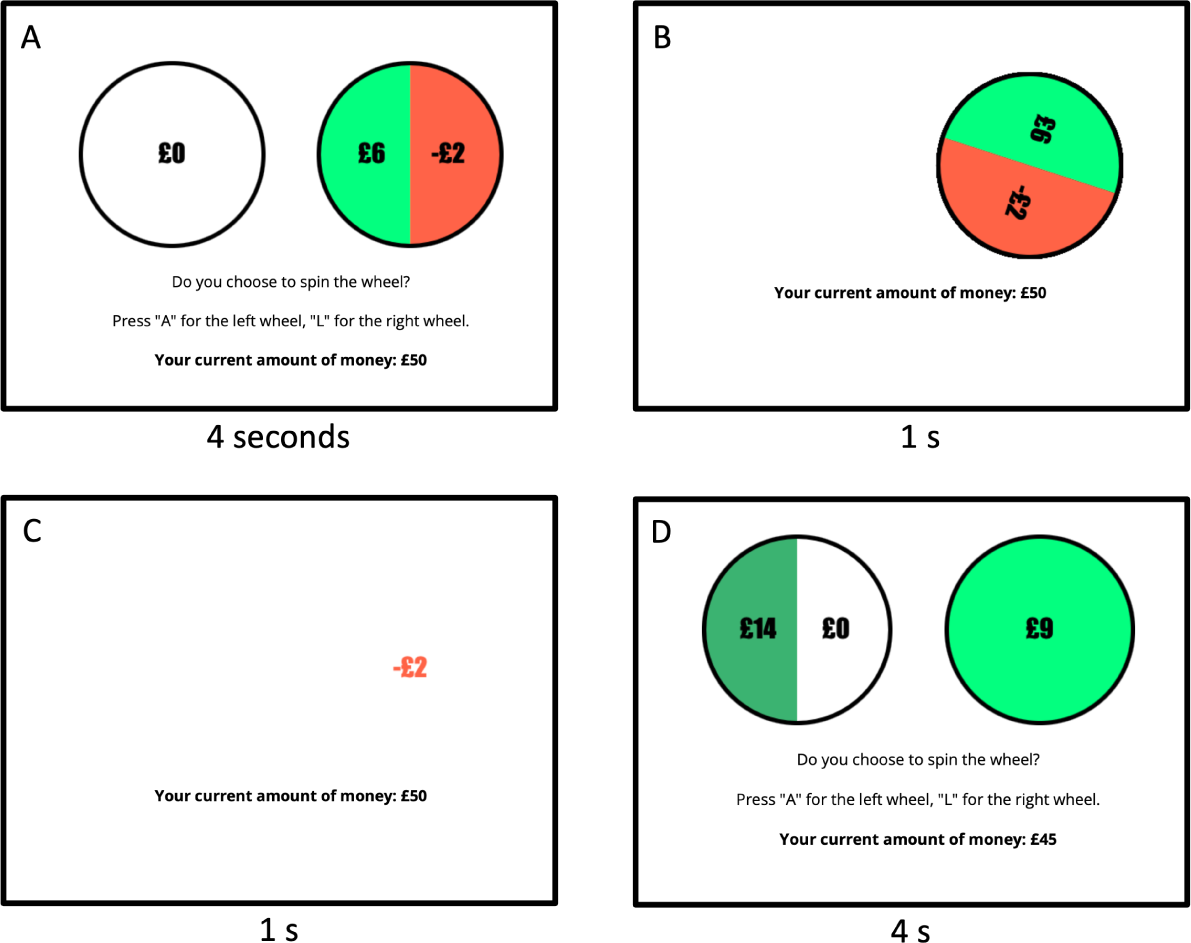
Task structure. (A) The frame shows a mixed gamble trial. The wheels are shown for 4s. If a participant does not make a choice within this time frame, the trial results in a time out. Once a wheel is chosen a 1s animation is shown (B), followed by the amount won/lost shown for 1s (C). The last frame (D) shows a gain-only trial, which has the same trial structure.

### 2.4 Questionnaires

At the end of the gambling task, participants were asked to complete several questionnaires. First, we administered the State Trait Anxiety Inventory (STAI) (Spielberger, 1977) to measure state and trait anxiety scores, two measures of anxiety that have widely been used in computational studies (Raymond, Steele, and Seriès, 2017), and the GAD-7 (Spitzer et al., 2006) questionnaire to assess Generalised Anxiety Disorder. Following these, participants were asked 13 questions about the COVID pandemic. Four questions asked about the subject vaccination status, the type of mask used throughout the pandemic, how anxious they were feeling at the beginning of the pandemic and how closely they followed government guidelines during the pandemic. We matched these questions with four questions asking about the subject’s future attitude towards vaccine boosters, masks, anxiety about attitude towards rules and guidelines in the case of a new surge of COVID cases. The last 5 questions asked about the subject’s general attitude towards COVID and vaccines. All these questions are rated on a Likert-scale of [1-5] and, based on the answers, we computed a COVID score where higher values corresponds to more cautious behaviour during the COVID pandemic (for example, completing the vaccination cycle, following government guidelines and wearing N95 masks). The COVID questionnaire and rating scale is reported in full in the Supp. Inf.

It can be seen from table 1 that no statistically significant difference were observed between our samples of vaccinated and unvaccinated individuals with respect to state and trait anxiety and GAD-7 scores. However, a statistically significant difference across COVID scores is present - even when the number of vaccination is not included in the score - showing that our questionnaire is able to pick up the different attitude towards COVID between unvaccinated and vaccinated individuals.

### 2.5 Prospect Theory Models

We fitted three different models derived from Prospect Theory to the subjects’ data. The models are the same as those used in Charpentier et al. (2017): the first model measures both risk and loss aversion, while the other two are nested models which only measure loss and risk respectively. The models have been fitted using both maximum likelihood estimation (MLE) in MATLAB and using hierarchical Bayesian methods implemented in the hBayesDM R library (Ahn, Haines, and Zhang, 2017).

The first model (allP) captures both risk and loss aversion and is described by the following equations

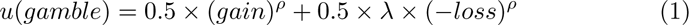

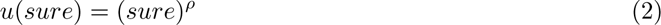

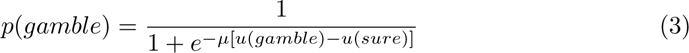

where *u* is the subjective utility of gambling or choosing the sure option, *gain* and *loss* are the numeric values shown on the gambling wheel and *sure* is the value of the sure option. The difference between the subjective utilities is then passed through a softmax function parameterised by inverse temperature *µ* (eq. (3)); this remains the same across all models. The model has three parameters:

- *λ* represents loss aversion. When *λ >* 1, the subject weights losses more than gains and it is therefore loss aversed; *λ <* 1 represents the inverse.
- *ρ* represents risk aversion. When *ρ <* 1, the subjects is risk aversed, whereas with *ρ >* 1 the subject is risk seeking.
- *µ* is the inverse temperature of the action selection function where higher values of *µ* represent less stochastic decisions.

The second model (noRA) has two parameters, one for loss aversion *λ* and one for the inverse temperature *µ*:

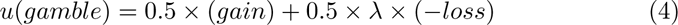

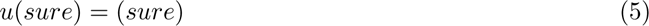

and the third model (noLA) similarly only includes a parameter for risk aversion *ρ* and the inverse temperature *µ*:

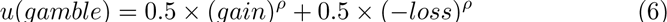

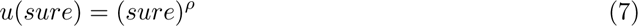

The Prospect Theory models were fitted using both maximum likelihood estimation (MLE) or hierarchical Bayesian (HB) methods. Maximum Likelihood models were fitted using MATLAB R2021b. We run constrained optimisation (fmincon with default settings) for *λ, ρ, µ ∈* [0*, ∞*]. For each subject we run optimisation 20 times and we select the best set of parameters according to their negative log likelihood. Hierarchical Bayesian models were fitted using R version 3.6.3 and hBayesDM (Ahn, Haines, and Zhang, 2017) version 1.2.1 which, in turn, uses Stan (Carpenter et al., 2017) version 2.21.7. We fitted 4 chains for each model, with 2000 burn-in samples and 6000 samples (8000 samples in total). We visually inspected traces for each parameter for good mixing and checked that the Gelman–Rubin statistic *R*^ (Gelman and Rubin, 1992) was close to 1 for all fitted parameters to test for convergence.

#### 2.5.1 Parameter Recovery

We performed extensive simulation and parameter recovery for the first model using both maximum likelihood estimation (MLE) and Hierarchical Bayesian methods (HB). We tested experimental designs comprising 148, 222 and 296 trials, with different combinations of Indifference Points and low/high anxiety parameters. For each combinations of parameters and Indifference Points(IP) we simulated 100 participants. Simulation parameters for low and high anxiety agents were taken from the existing literature (Charpentier et al., 2017). We report correlations using the more robust Spearman corerlation coefficient, due to outliers resulting from the MLE fitting procedure. Across all combinations of parameters, fitting procedure and number of trials, recovery is successfull, with at least a Spearman *ρ >* 0.64. Parameter recovery results for 148, 222 and 296 trials and full details of the parameter recovery set up are reported in the Supp. Inf. We set the length of the experiment to 148 trials due to the similar results of the parameter recovery across different number of trials.

#### 2.5.2 Model Selection

We fitted all three models using both Maximum Likelihood Estimation and Hirarchical Bayesian methods. We compared them using the Akaike Information Criterion (AIC) and the Bayesian Information Criterion (BIC) for the parameters fitted using MLE and we compared them using Leave One Out Information Criterion (LOOIC) for the HB fitting.

We then fit the winning model with six different hierarchical priors specifications on the parameters (similarly to Ahn, Vasilev, et al., 2014; Aylward et al., 2019): a single prior for each parameter encompassing all participants, two separate priors for each parameter for low and high anxiety subjects according to the GAD-7 median split, two separate priors for each parameter for low and high anxiety subjects according to the STAI Trait median split, two separate priors for each parameter for vaccinated and unvaccinated subjects, four different priors for each parameter for the combinations of GAD-7 low/high anxiety and vaccinated/unvaccinated subjects and four different priors for each parameter for the combinations of STAI Trait low/high anxiety and vaccinated/unvaccinated subjects. Throughout the rest of this work we refer to these models as single prior, GAD-7/Trait anxiety priors, vaccination priors and GAD-7/Trait anxiety and vaccination priors for convenience.

### 2.6 Hierarchical Drift Diffusion Models

Hierarchical Drift Diffusion Models (HDDMs) are a type of sequential sampling models which allow to discriminate between the different elements that are involved when making a decision. Contrary to the Prospect Theory models, HDDMs model both choice and reaction times concurrently. Our analysis focused on four main drift diffusion models components: the boundary separation a, the non-decision time t, the drift-rate v and the starting point z. We implement and fit the models using the HDDM library version 0.8.0 (Wiecki, Sofer, and Frank, 2013) and Python 3.6.15. Parameters in HDDM are allowed to “depend on” certain conditions, meaning that the model will estimates separate parameters for each condition. For this analysis, our conditions correspond to the type of trials, i.e., mixed-gamble vs. gain-only, and we design models in which all possible combinations of parameters are varying across types of trials, with the other ones being the same across conditions. In total we test 16 models, namely: none, a, t, v, z, at, av, az, tv, tz, vz, atv, atz, avz, tvz, atvz. Models are separately fitted the low and high anxiety groups based on the GAD-7 median score, similarly to the Prospect Theory models described above.

Each model was fitted using HDDM’s MCMC sampler with 4 chains, 1000 burn in samples and 3000 samples. We visually inspected the MCMCM traces for good mixing and the autocorrelation plots for low autocorrelations, we also checked the Gelman-Rubin *R*^ statistics for convergence (0.9 *< R*^ *<* 1.1). Models were compared using the Deviance Information Criterion (DIC) (Ando, 2007) provided by the HDDM library.

#### 2.6.1 Parameter Recovery

We conducted parameter recovery for the HDDMs by simulating 115 agents (high anxiety: 54, low anxiety: 61) using the atvz model. Simulation parameters were sampled from Gaussian distributions with mean and standard deviation coming from the atvz model estimates on our dataset. The full parameter recovery results are available in the Supp. Inf., where we obtained good recoverability for the non-decision time t (*r ∈* [0.65, 0.89]), mixed recoverability for the drift rate v (*r ∈* [0.34, 0.55]) and poor recoverability for starting point z and boundary separation a (*r ∈* [0.06*−*0.5] likely due to the model fitting procedure introducing correlations between the parameters (see below).

### 2.7 Model Agnostic Analysis

We report differences in parameters fitted using hierarchical Bayesian methods using the difference between the 95% Highest Density Interval (HDI) of the two groups estimates. If the resulting HDI did not encompass zero, we conclude that there is a significant difference between the two groups. All *t*-tests are two-sided with a significance level of *α* = 0.05. All correlation coefficients are Pearson’s *r*, unless stated otherwise. Bayesian *t*-tests and correlations have been carried out using JASP (JASP Team, 2023) version 0.17.0. For multiple comparisons tests, we report ‘native’ *p*-values together with the Bonferroni corrected *p*-values. We applied Bonferroni correction for each family of tests we performed: 18 tests related to reaction times (corrected *α* = 0.0027) and 8 tests related to individual choice data (corrected *α* = 0.00625).

## 3 Results

### Choice Data Analysis

There were no group differences across percentages of gamble across either the low/high anxiety groups or the vaccinated/unvaccinated groups (all *p >* 0.144; see Supp. Inf.). Percentage of gambles across mixed gamble and gain-only trials are not significantly correlated (*r*(115) = 0.148, *p* = 0.116, BF_10_ = 0.396) indicating that subjects must use different strategies in these different types of trials.

Participants were payed a bonus up to £1, proportional to their final amount of money collected at the end of the gambling trials. The final bonus won by participants was on average £0.37 (£0.08) with no significant difference between vaccination status (vaccinated: £0.369 (£0.082); unvaccinated £0.371 (£0.077); *t*_113_ = 0.131, *p* = 0.896) or low/high anxiety (low anxiety: £0.318 (£0.087); high anxiety £0.359 (£0.070); *t*_113_ = 1.496, *p* = 0.137).

### Risk and Loss Aversion

The Hierarchical Bayesian (HB) fitting method produced more stable results compared to Maximum Likelihood Estimation (MLE) during both parameter recovery and when fitting the participants’ data. Model 1 is the best model according to both the AIC, BIC and LOOIC scores and it is the model used for the rest of this analysis (full results in Supp. Inf.).

Our data is best fitted using a single prior for all participants (table 2), with the model fitted using the GAD-7 median split priors being a close second.

**Table 2:** Model 1 LOOIC scores. Model 1 LOOIC scores for different priors specifications.

|  | LOOIC |
| --- | --- |
| Single prior | **13065.12** |
| GAD-7 Anxiety priors | 13068.45 |
| STAI Trait Anxiety priors | 13086.08 |
| Vaccination priors | 13083.36 |
| GAD-7 Anxiety and vaccination priors | 13101.54 |
| STAI Trait Anxiety and vaccination priors | 13086.81 |

Due to the negligible difference in LOOIC scores between the single prior model and the GAD-7 anxiety prior model the rest of the analysis will report results from the GAD-7 anxiety priors model. This decision is also backed up by previous research (Valton, Wise, and Robinson, 2020) which recommends to account for hypothesised group differences in computational modelling. For completeness, we report results for the single prior model in the Supp. Inf.

We report the fitted parameters for the GAD-7 anxiety priors model in table 3 and in fig. 2(A-C). Consistently with the Prospect Theory literature (Kahneman and Tversky, 1979), individuals are on average risk averse (*ρ <* 1) and they value losses twice as much as wins (*λ ≈* 2). The high anxiety group shows increased loss aversion (BF_10_ = 2.156) and reduced inverse temperature (BF_10_ = 2.407) compared to the low anxiety group. No difference in risk aversion (BF_10_ = 0.239) was found between the low and high anxiety groups. GAD-7 scores are significantly correlated with loss aversion (*r*_115_ = 0.193, *p* = 0.039, BF_10_ = 0.953, fig. 2 (D)) and inverse temperature (*r*_115_ = *−*0.196, *p* = 0.035, BF_10_ = 1.033), but not risk aversion (*r*_115_ = 0.075, *p* = 0.429, BF_10_ = 0.159).

**Figure 2:**
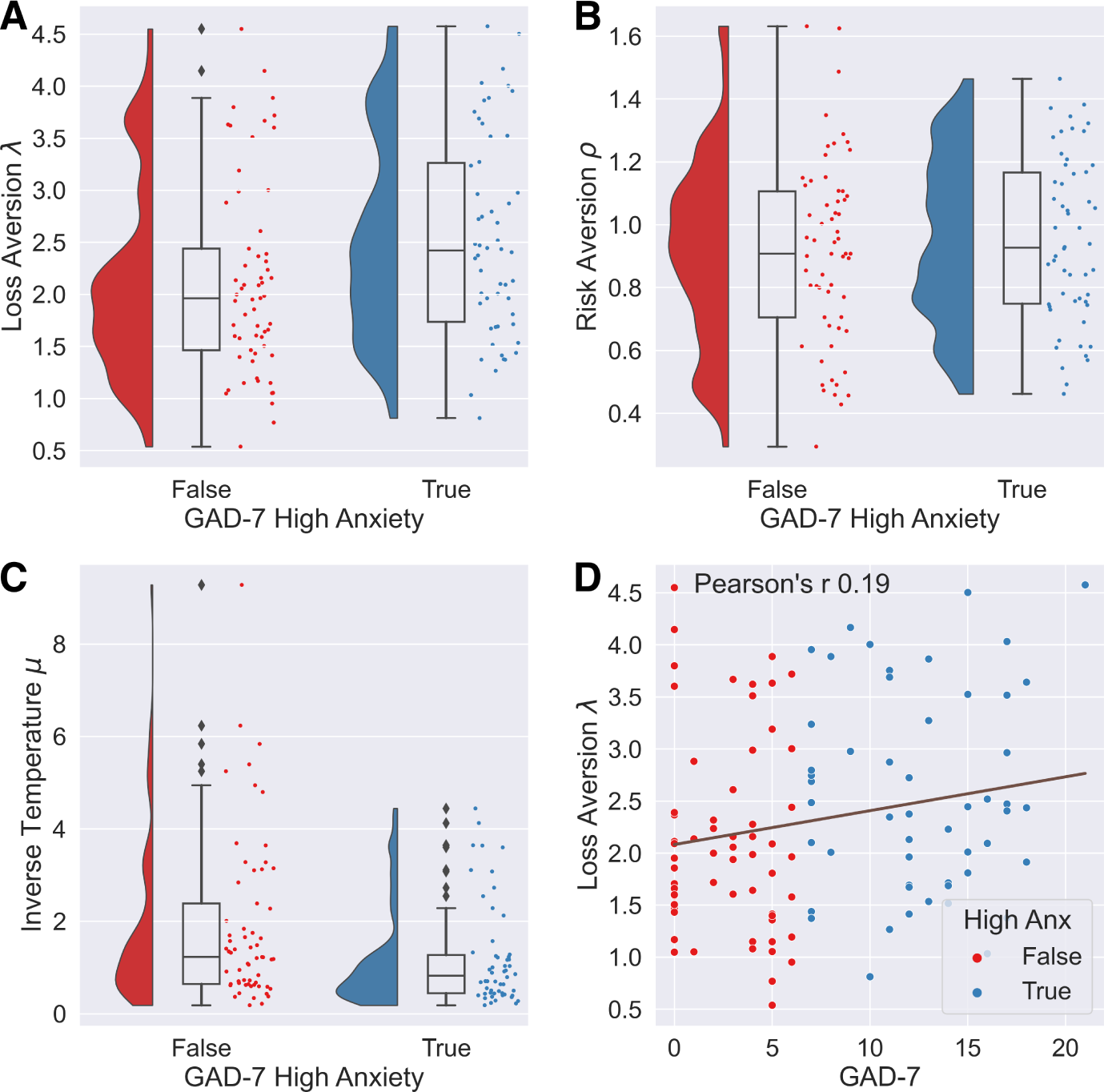
Model 1 fitted parameters. (**A-C**) Parameters distributions for the anxiety prior model. (**D**) Pearson correlation between GAD-7 scores and Loss Aversion parameter.

**Table 3:**
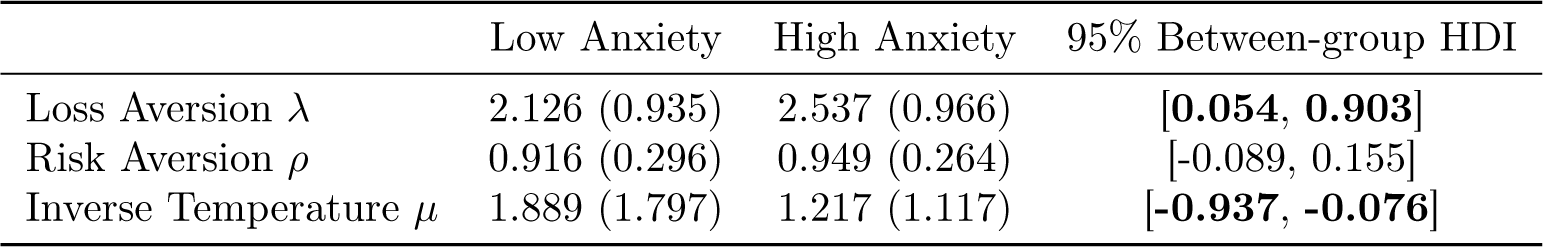
Model 1 fitted parameters. Mean (standard deviation) of the fitted parameters of Model 1 using the anxiety prior based on the GAD-7 scores median split. Parameters for which the between-group 95% HDI does not encompass zero (in bold) are considered different across groups.

We fitted a logistic regression model with low/high anxiety (based on GAD-7 median split as above) as the dependent variable and loss *λ* and risk *ρ* aversion parameters as independent variables. We find that loss aversion was a statistically significant predictor of anxiety group (*z* = 2.242, *p* = 0.025) while risk aversion was not (*z* = 0.641, *p* = 0.522).

Despite previous research pointing toward differences between low/high STAI Trait anxiety scores, our model fails to capture any significant group difference in its parameters. This is surprising given a strong correlation between the STAI Trait Anxiety scores and the GAD-7 scores (*r*_115_ = 0.830, *p <* 0.001, BF_10_ *>* 10^27^).

### COVID Score analysis

Our data do not support any correlation between fitted parameters and COVID score (or the COVID-Past/Future/Generic scores). We report a positive correlation between State Anxiety and COVID score (*r*(115) = 0.194, *p* = 0.037, BF_10_ = 0.993), State Anxiety and COVID-Future score (*r*(115) = 0.193, *p* = 0.038, BF_10_ = 0.972) and State Anxiety and COVID-Generic score (*r*(115) = 0.199, *p* = 0.032, BF_10_ = 1.113). We also find a near significant correlation between State Anxiety and COVID-Past score (*r*(115) = 0.159, *p* = 0.089, BF_10_ = 0.484)

### Reaction Times

Mean reaction times did not differ across low and high anxiety subjects (low anxiety: 1.254 (0.470) seconds; high anxiety: 1.213 (0.438) s; *t*_113_ = *−*0.791, *p* = 0.430) or vaccinated and unvaccinated (vaccinated: 1.208 (0.439) s; unvaccinated: 1.264 (0.472) s; *t*_113_ = *−*1.105, *p* = 0.271). We observe a statistically significant difference in reaction times between low/high anxiety subjects in mixed gamble trials in which participants decided not to gamble: the high anxiety group chooses not to gamble faster compared to the low anxiety group (low anxiety: 1.384 (0.391) s; high anxiety 1.240 (0.245) s; *t*_113_ = *−*2.326, *p* = 0.021, not significant under a Bonferroni correction with *α* = 0.0027).

### Hierarchical Drift Diffusion Models

All models showed good mixing, low autocorrelation and Gelman-Rubin statistics within the desired bounds. We find that the atvz model is the best fitting model according to DIC scores (see Supp. Inf.). The winning atvz model produced separate posterior distributions on mixed-gamble and gain-only trials for each parameter. We report the between-group *t*-tests and 95% HDI difference of these distributions in table 4.

**Table 4:** HDDM group differences. Group differences between the low and high anxiety groups in the atvz model. a is boundary separation, t is non-decision time, v is drift-rate, z is starting point.

|  | t-tests (p-values) |  | 95% HDI |  |
| --- | --- | --- | --- | --- |
|  | mixed-gamble | gain-only | mixed-gamble | gain-only |
| **a** | 0.051 | 0.604 | [-0.168, 0.001] | [-0.085, 0.102] |
| **t** | 0.470 | 0.101 | [-0.100, 0.042] | [-0.135, 0.011] |
| **v** | 0.160 | 0.053 | [-0.410, 0.072] | [-0.048, 0.452] |
| **z** | 0.456 | 0.081 | [-0.033, 0.021] | [-0.048, 0.011] |

No significant difference can be found between groups for any of the parameter, with the exception of a near significant difference in boundary separation a in mixed-gamble trials and its 95% HDI difference very close to not encompass zero. High anxiety subjects show reduced boundary separation (high anxiety ā = 1.514; low anxiety ā = 1.596) in mixed-gamble trials, a result which is difficult to interpret in terms of risk and loss aversion when presented on its own. In fact, the boundary separation a is is significantly correlated with the drift rate v (*r*_115_ = 0.403, *p <* 0.001), which is in turn significantly correlated with the starting point z (*r*_115_ = 0.330, *p <* 0.001), making the overall interpretability of these results quite difficult. Furthermore, parameter recovery for this model introduced correlations between parameters (see Supp. Inf.) which made for poor recoverability.

We investigate whether having simpler models, e.g. with fewer parameters, may provide better interpretability of the results and lower correlations between the parameters. We repeat the analysis on the subsequent 5 best models, in order: atv, tvz, tv, atz and tz.

Models atv, tvz, tv failed to produce significant differences across parameters, whilst also having significant correlations across their parameters (more results are available in the Supp. Inf.).

Model atz shows a statistically significant difference between low and high anxious groups in starting point z in mixed-gamble trials (*t*_113_ = *−*2.000, *p* = 0.047) and the 95% HDI for the same parameter is very close to not encompass zero ([*−*0.059, 0.001]). The starting point for the more anxious subjects is closer to 0.5 (mean = 0.51), whereas the low anxiety group shows a bias towards the gambling option (mean = 0.54), which is consistent with the increased loss aversion for the low anxiety group found using the Prospect Theory models. This model also shows a weak, close to significance, correlation between boundary separation and starting point in mixed-gamble trials (*r*_115_ = 0.175, *p* = 0.061). Parameters obtained for the gain-only trials are all correlated to each other: a and t (*r*_115_ = 0.196, *p* = 0.035), a and z (*r*_115_ = *−*0.205, *p* = 0.027) and t and z (*r*_115_ = *−*0.184, *p* = 0.047)

Model tz also shows a statistically significant difference between low and high anxious groups in starting point z in mixed-gamble trials (*t*_113_ = *−*2.071, *p* = 0.040) and the 95% HDI for the same parameter is very close to not encompass zero ([*−*0.057, 0.002]). The starting point for the more anxious subjects is again closer to 0.5 (mean = 0.51), whereas the low anxiety group shows a bias towards the gambling option (mean = 0.54). In this model, non-decision time t and starting point z are not correlated in mixed-gamble trials (*r*_115_ = 0.004, *p* = 0.961), and close to significantly correlated in gain-only (*r*_115_ = *−*0.161, *p* = 0.085). Starting points are, close to significance, negatively correlated with GAD-7 scores (*r*_115_ = *−*0.182, *p* = 0.051). Starting points are also strongly negatively correlated with the loss aversion parameters (*r*_115_ = *−*0.690, *p <* 0.001) and weakly correlated with the risk aversion parameters (*r*_115_ = 0.212, *p* = 0.022).

Despite the lower number of parameters of models atz and tz, parameter recovery still presents low recoverability for a (*r ∈* [0.14, 0.53] for the atz model) and z (*r ∈* [0.28, 0.62] for atz; *r ∈* [0.13 *−* 0.54] for tz). Parameter recovery also introduced correlations between the recovered parameters (see Supp. Inf. for full results of the parameter recovery).

## 4 Discussion

In this study we found that individuals with high levels of Generalised Anxiety Disorder show increased loss aversion compared to individuals with lower anxiety levels. No differences in risk aversion were found between the low and high anxiety groups. Our results are consistent with a recent study by Xu et al. (2020), which found increased levels of loss aversion and no differences in risk aversion in individuals with high trait anxiety. Xu et al. used a Prospect Theory model similar to the one used in our study but their task design differs in that they only employed mixed gamble trials in which expected value and variance of the gain/loss amounts were manipulated. Our task, on the other hand, like the one used by Charpentier et al. (Charpentier et al., 2017), present different types of trials (mixed gamble and gain-only). Notably, our results do not replicate Charpentier et al.’s differences in risk aversion and instead show differences in loss aversion, where they did not find any. This suggests that more work is needed in this field, both to test the robustness of these results and understand the differences observed across different studies. Several competing factors can explain these mixed results. First, our study and Xu et al.’s study were conducted on subclinical populations, whereas Charpentier et al.’s study was conducted on a clinical population, which may indicate possible behavioural differences between subclinical and clinical populations of anxiety. Charpentier et al.’s study also included an emotional manipulation task which could have had an impact on decision-making that may not be picked up by the models used. Second, Xu et al.’s and Charpentier et al.’s studies employed STAI Trait anxiety scores as a measure of anxiety, whereas in our study results only hold for GAD-7 anxiety scores and not for STAI Trait anxiety scores. This may highlight the use of largely overlapping but different cognitive mechanisms captured by STAI Trait anxiety scores and GAD-7 scores, despite these being highly correlated both in our study (*r*_115_ = 0.83) and more generally in the literature (Doi et al., 2018). Third, our experiment explicitly instructs participants to maximise their final rewards in order to win a larger bonus payment while participants are able to see the result of each gamble (and therefore the amount of money won or lost) at each trial. In Xu et al.’s experiment, participants were only shown the result of a trial selected at random, with the won/lost amount being added/removed from a fixed amount of money received by the participants before the experiment; in Charpentier et al.’s experiment, participants were shown the results of 10 trials selected at random which were then averaged and either added or removed from a fixed amount of money to reward the participant. This difference in experimental design may explain the overall high values (i.e., reduced risk aversion) that our model fitted for the risk aversion *ρ* parameter (overall: 0.931 (0.281), low anxiety: 0.916 (0.296), high anxiety: 0.949 (0.264)), compared, for example to a lower mean value of *ρ* reported by Charpentier et al. (overall: 0.713 (0.458), low anxiety: 0.875 (0.537), high anxiety: 0.564 (0.313)). Our study’s explicit instructions to maximise rewards could therefore explain our values of *ρ* close to 1, participants may be motivated to accept more gambles than they normally would in order to increase their in-game scores and their proportional bonus payments. During the experiment’s practice trials, we adjust the trials expected values to each participant’s sensibility, and we remove participants for which this procedure was not successful. As a result of this, the average expected value of all trials was positive for each participants. Therefore, increased gambling would lead to net positive wins.

We did not find statistically significant differences in reaction times across anxiety levels, apart from the high anxiety group being faster at choosing not to gamble (i.e., avoiding the chance of losing money) in mixed gamble trials. This could be explain in terms of reduced boundary separation, shorter non-decision time, higher drift rate, less biased starting point or a combination of these. The results from the best fitting HDDM model atvz partially support a difference in boundary separation a (p = 0.051, 95% HDI [-0.168, 0.001]) in mixed gamble trials, with more anxious subjects having smaller boundary separations This could in part explain the faster reaction times shown by anxious subjects when rejecting a gamble in mixed-gamble trials. But, the correlations between the parameters of this model (and the next 3 best fitting ones) make it difficult to interpret these results, as these may be due to the parameters trading off against each other rather than actual differences. Models atz and tz, respectively the 5th and 6th best fitting model according to DIC scores, both report significant differences in starting point z in mixed-gamble trials, with weak correlation between starting point and boundary separation in the atz model. Both models found lower, less biased starting points in mixed-gamble trials for the more anxious subjects. This result can be interpreted in terms of increased loss aversion, as subjects are less biased towards the gambling option in the trials with the chance of loosing money. Model tz also shows a near to significance correlation between less biased starting points and GAD-7 scores. Despite the lack of definitive results, reduced boundary separations and less biased starting points in mixed-gamble trials for high anxious subjects seem to be a common theme across the best fitting models. Our results, faster reaction times when rejecting a gamble in mixed-gamble trials and a strong influence of starting point bias in the high anxiety group, are consistent with the pattern of results reported by Zhao, Walasek, and Bhatia (2020) for healthy subjects. Further research should therefore focus on understanding whether anxiety exacerbate these factors. A limitation of this study comes from the chosen trials structure - the original trials structure by Charpentier et al. (2017) was intentionally kept the same in an effort to reproduce the original Prospect Theory results. Exploratory analysis conducted on this dataset using traditional DDMs found that the task configuration (98 mixed-gamble trials, 50 gain-only trials) resulted in poor parameter recovery. Therefore, we believe that further experiments should test HDDMs models on tasks specifically designed for them; by, for example, performing extensive parameter recovery and simulations during the task design stage.

We also highlight how hierarchical Bayesian methods provide better and more reliable fitted parameters compared to Maximum Likelihood estimation, as measured in terms of better parameter recovery.

Previous questionnaires-based research (Millroth and Frey, 2021) has highlighted how anxiety and negative individual disposition towards risk and uncertainty have an important role in predicting strong fear responses related to the COVID-19 pandemic. However, in our study, the hypothesis that individual levels of risk and loss aversion could be linked to decision-making in the context of the COVID pandemic was rejected. Simple economic decision-making tasks like the one we employed might not be able to capture the intrinsic complexity of decision-making in a multifaceted situation like the COVID pandemic in which economic, health and social factors are all at interplay and more complex naturalistic tasks (Schonberg, Fox, and Poldrack, 2011) may be required to better study these relationships. A recent review about predictors of vaccine hesitancy (Hudson and Montelpare, 2021) includes risk aversion as one of the possible individual difference factors which influence vaccine uptake. Risk aversion could impact vaccine hesitancy in two different and opposite ways. A risk averse individual might value the unknown side effects of a vaccine as riskier than the effects of a known disease and would only get vaccinated due to work or travel requirements. At the same time, an individual might find the dangers posed by a vaccine-preventable disease as riskier than the possible side effects of a vaccine. Loss aversion may also contribute to decisions taken during the pandemic in different ways. For example, workers who would received little protection in case of loss of income due to self isolation following a COVID infection may value the loss of their working hours as more important than the (individual and public) health gains given by self isolating in case of mild COVID symptoms. Therefore, assuming vaccinations and stronger adherence to healthcare guidelines as the risk averse behaviours is a limitation in our study, as some individuals may consider these as more risk- and/or loss-prone actions. Hence, our self-report COVID questionnaire should be expanded to include questions designed to account between these different views.

In conclusion, our results indicate that subjects with high levels of Generalised Anxiety Disorder show increased loss aversion, and no differences in risk aversion, as measured by computational parameters fitted on data from a economic decision-making task. Future computational studies should focus on disentangling the effects of motivation and reward on risky decision-making and to better study the temporal patterns that might affect decision-making. These findings may help clinicians in tailoring behavioural interventions in order to overcome decision biases. A previous study (Lorian, Titov, and Grisham, 2012) provides evidence for a reduction of risk-averse behaviours following a cognitive behavioural therapy treatment. Therefore, future work could explore the benefits of including loss aversion as a treatment outcome in similar intervention.

## 5 Data Accessibility Statement

The code and data of this study are available on the Open Science Framework (OSF) at https://osf.io/xqwr5/.

## 8 Authors’ Contributions

- F.F.: Conceptualisation, Methodology, Software, Investigation, Formal analysis, Visualisation, Writing - Original Draft, Writing - Review & Editing.
- J.A.: Conceptualisation, Software, Writing - Review & Editing.
- P.S.: Conceptualisation, Writing - Review & Editing, Supervision, Funding acquisition

## A COVID Questionnaire

All items were scored between 1 to 5, we report the questions and the different answers within square brackets. The dots (*•*) indicate the questions about past behaviours, the asterisks (*) indicate the questions about future behaviours and the dashes (-) indicate the questions about general attitude towards COVID. We group these questions in COVID-Past (*•*), COVID-Future (*) and COVID-General (-) scores.

• How many COVID vaccinations have you had? [0, 1, 2, 3, 4+]
* Would you get another booster dose if offered? [Very Unlikely, …, Very Likely]
• What type of mask did you usually wear throughout the pandemic? [No Mask, Scarf/Bandana/Other, Reusable Fabric Mask, Surgical/Disposable Mask, Respirator (N99/N95)]
* What type of mask would you wear more often in case of a new surge in COVID cases? [No Mask, Scarf/Bandana/Other, Reusable Fabric Mask, Surgical/Disposable Mask, Respirator (N99/N95)]
• How anxious were you about coronavirus at the beginning of the pandemic in March 2020? [Not at all Anxious, …, Extremely Anxious]
* How anxious are you about a new increase in COVID cases in the upcoming Winter? [Not at all Anxious, …, Extremely Anxious]
• How closely have you followed the government rules and guidelines during the pandemic? [I have not followed any of the rules, …, I have followed all of the rules]
* In case of a hypothetical new pandemic how closely would you follow new government rules and guidelines? [I would not follow any of the rules, …, I would follow all of the rules]
- I think Coronavirus vaccinations should be mandatory. [Strongly Disagree, …, Strongly Agree]
- During the pandemic I wore a mask and social distanced as often as I could. [Strongly Disagree, …, Strongly Agree]
- I think that Coronavirus restrictions introduced in 2020 were necessary and important. [Strongly Disagree, …, Strongly Agree]
- I am worried or have been worried about coronavirus affecting my health. [Strongly Disagree, …, Strongly Agree]
- I am worried or have been worried about coronavirus affecting the health of my friends and family. [Strongly Disagree, …, Strongly Agree]

## B Exclusion Criteria

Different types of exclusion criteria have been used. During the gambling part of the experiment, 4 trials were used as attention checks. In these trials the subjects had to choose between a win of £0 and a loss in the range £[-30,-5] together with a sure option of £0. We discarded participants who chose the gambling option for 2 or more of these 4 attention checks. Another attention check was included in the questionnaire part of the experiment. We removed 22 participants who failed 2 or more gambling attention checks whereas no participant failed the questionnaire attention check. Four participants were removed due to incomplete demographic information.

We also removed participants who did not show any differences in percentage of gambling as a function of the trials’ expected values. We remove participants which have a difference in average percentage of gamble lower than 5% between the upper half of expected values and the lower half of expected values. Participants should be able to adapt their gambling to the different expected values of the trials (i.e., higher expected values should lead to higher percentage of gambles). An example of this can be seen in fig. 3. We removed 22 participants using this criteria.

**Figure 3:**
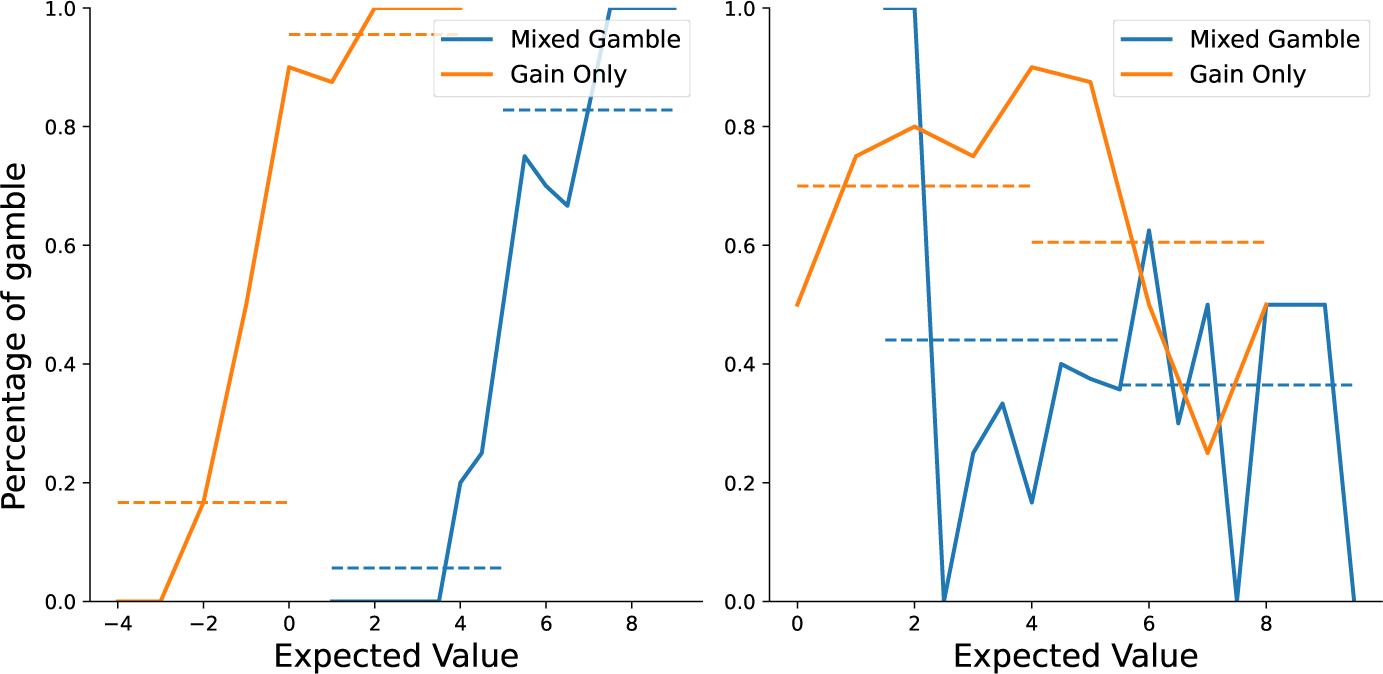
On the left, a participants which correctly adapted their choices to the expected value of the trials. On the right, a participant which incorrectly reduces their percentage of gambles for gambles with higher expected values. The participant on the right has been removed from further analysis.

In total we conducted the analysis on 115 participants.

## C Parameter Recovery and Model Comparison

Parameter recovery was conducted simulating 100 subjects for each combination of parameters/indifference points. Parameters were sampled from Gaussian distributions using the fitted parameters reported by Charpentier et al. (2017) (reported here in table 5). Indifference points were sampled from Gaussian distributions with SD = 2 and means as follows: High IP mixed gamble = 8, High IP gain only = 4, Low IP mixed gamble = 2, Low IP gain only = 1.

**Table 5:**
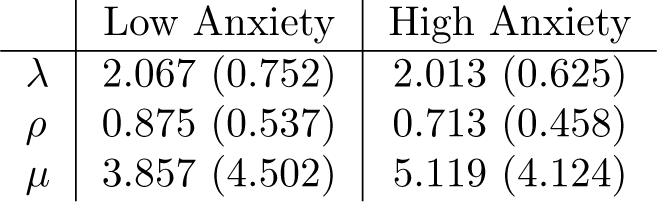
Parameters from Charpentier et al. (2017) used for the parameter recovery. Mean (Standard deviation).

The results for the parameter recovery are reported in table 6. Due to outliers in the MLE fitting of the data we report the more robust Spearman correlation coefficients.

**Table 6:** Parameter recovery for 148, 222 and 296 trials. We report Spearman Correlation Coefficients. IP = Indifference Point.

|  | MLE |  |  | HB |  |  |
| --- | --- | --- | --- | --- | --- | --- |
| | $\lambda$ | $\rho$ | $\mu$ | $\lambda$ | $\rho$ | $\mu$ |
| 148 trials |  |  |  |  |  |  |
| High Anxiety / Low IP | 0.82 | 0.88 | 0.88 | 0.91 | 0.96 | 0.90 |
| High Anxiety / High IP | 0.71 | 0.93 | 0.87 | 0.92 | 0.96 | 0.84 |
| Low Anxiety / Low IP | 0.80 | 0.88 | 0.91 | 0.95 | 0.97 | 0.79 |
| Low Anxiety / High IP | 0.84 | 0.90 | 0.84 | 0.89 | 0.94 | 0.75 |
| 222 trials |  |  |  |  |  |  |
| High Anxiety / Low IP | 0.82 | 0.87 | 0.95 | 0.93 | 0.99 | 0.94 |
| High Anxiety / High IP | 0.77 | 0.91 | 0.78 | 0.89 | 0.97 | 0.92 |
| Low Anxiety / Low IP | 0.77 | 0.90 | 0.93 | 0.95 | 0.98 | 0.92 |
| Low Anxiety / High IP | 0.64 | 0.87 | 0.90 | 0.87 | 0.95 | 0.74 |
| 296 trials |  |  |  |  |  |  |
| High Anxiety / Low IP | 0.85 | 0.97 | 0.94 | 0.94 | 0.97 | 0.97 |
| High Anxiety / High IP | 0.82 | 0.96 | 0.87 | 0.95 | 0.98 | 0.93 |
| Low Anxiety / Low IP | 0.81 | 0.93 | 0.89 | 0.98 | 0.99 | 0.90 |
| Low Anxiety / High IP | 0.89 | 0.92 | 0.87 | 0.86 | 0.96 | 0.82 |

**Table 7:** AIC, BIC and LOOIC scores for the three models. Lower is better.

|  | MLE |  | HB |
| --- | --- | --- | --- |
|  | AIC | BIC | LOOIC |
| Model 1 (allP) | **46084.59** | **13768.63** | **13065.12** |
| Model 2 (noRA) | 54758.99 | 21868.35 | 14234.04 |
| Model 3 (noLA) | 52286.11 | 19395.47 | 18819.73 |

## D Choice Data Analysis

**Table 8:** Probability of gamble during trials. Here p(gamble-mixed) and p(gamble-gain) refers to the probability of gambling in, respectively, mixed gamble and gain only trials.

| | $p(\text{gamble})$ | $p(\text{gamble-mixed})$ | $p(\text{gamble-gain})$ |
| --- | --- | --- | --- |
| All participants | 0.587 | 0.576 | 0.609 |
| Low Anxiety | 0.597 | 0.603 | 0.586 |
| High Anxiety | 0.577 | 0.546 | 0.636 |
| Vaccinated | 0.578 | 0.553 | 0.628 |
| Unvaccinated | 0.597 | 0.601 | 0.589 |

## E Single Prior Model

Across all participants, the mean (and standard deviation) of the loss aversion parameter *λ* is 2.323(0.975), the risk aversion parameter *ρ* is 0.933(0.283) and the inverse temperature *µ* is 1.568(1.526). We report the fitted parameters for this model in table 9 and in fig. 4.

**Figure 4:**
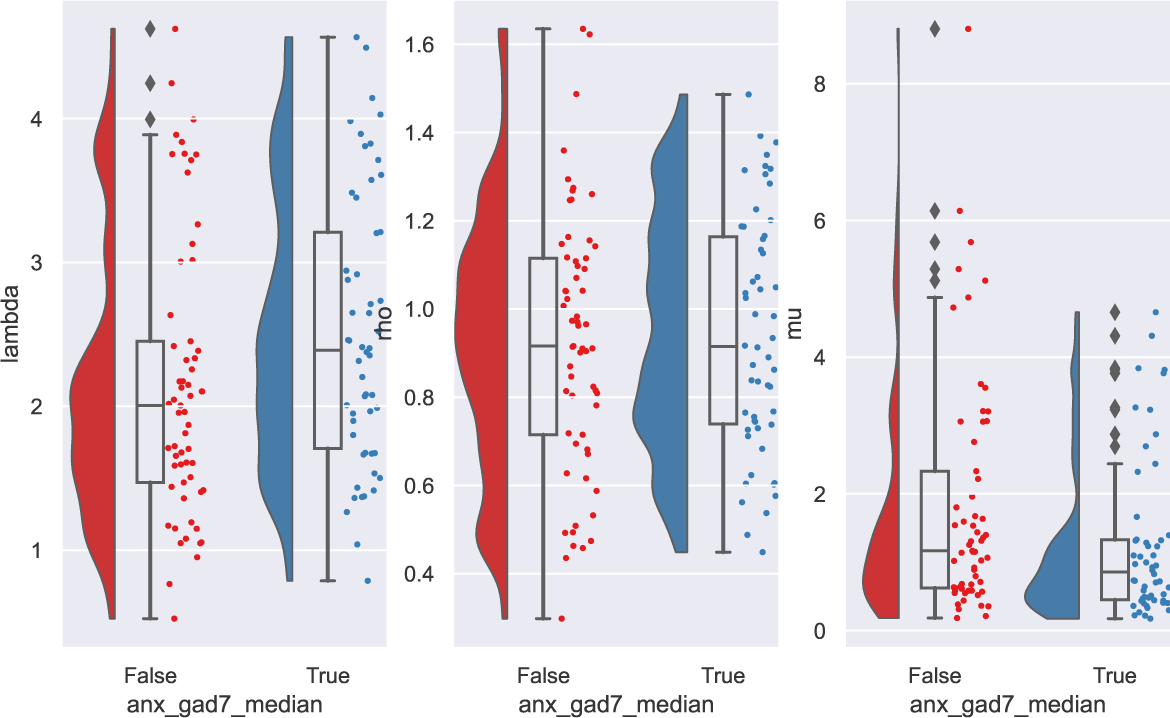
Parameters distributions for the single prior model.

**Table 9:** Mean (standard deviation) of the fitted parameters of Model 1 using a single prior (top table) and a GAD-7 score median split prior.

| Single prior | Loss Aversion $\lambda$ | Risk Aversion $\rho$ | Inverse Temperature $\mu$ |
| --- | --- | --- | --- |
| All participants | 2.323 (0.975) | 0.933 (0.283) | 1.568 (1.526) |
| Low Anxiety | 2.156 (0.971) | 0.924 (0.296) | 1.829 (1.748) |
| High Anxiety | 2.510 (0.955) | 0.943 (0.270) | 1.274 (1.178) |

| Anxiety prior | Loss Aversion $\lambda$ | Risk Aversion $\rho$ | Inverse Temperature $\mu$ |
| --- | --- | --- | --- |
| All participants | 2.318 (0.967) | 0.931 (0.280) | 1.573 (1.546) |
| Low Anxiety | 2.125 (0.935) | 0.915 (0.296) | 1.889 (1.797) |
| High Anxiety | 2.537 (0.966) | 0.949 (0.264) | 1.217 (1.116) |

We find a near significant group difference between the low and high anxiety groups in the single prior model in loss aversion *λ* (*t*_113_ = 1.964, *p* = 0.051), with the more anxious subjects showing higher levels of loss aversion, and in inverse temperature *µ* (*t*_113_ = *−*1.970, *p* = 0.051), with the more anxious subjects having lower values of *µ*. In the model fitted using the anxiety priors we find the same group differences, but now statistically significant (*λ* : *t*_113_ = 2.317, *p* = 0.022; *µ* : *t*_113_ = *−*2.371, *p* = 0.019). There was no difference in risk aversion across the two groups in either models (single prior *ρ* : *t*_113_ = 0.348, *p* = 0.727; anxiety prior *ρ* : *t*_113_ = 0.639, *p* = 0.523).

Risk and loss aversion are not correlated across individuals (*r*(115) = 0.004, *p* = 0.962), while inverse temperature is negatively correlated with both risk (*r*(115) = *−*0.624*, p <* 0.001) and loss aversion (*r*(115) = *−*0.333*, p <* 0.001). In the single prior model, GAD-7 scores are near significantly correlated to loss aversion (*r*_115_ = 0.168, *p* = 0.072) and inverse temperature (*r*_115_ = *−*0.166, *p* = 0.075). In the anxiety prior model, GAD-7 scores are significantly correlated with loss aversion (*r*_115_ = 0.192, *p* = 0.039) and inverse temperature (*r*_115_ = *−*0.196, *p* = 0.035).

## F Model 1

**Figure 5:**
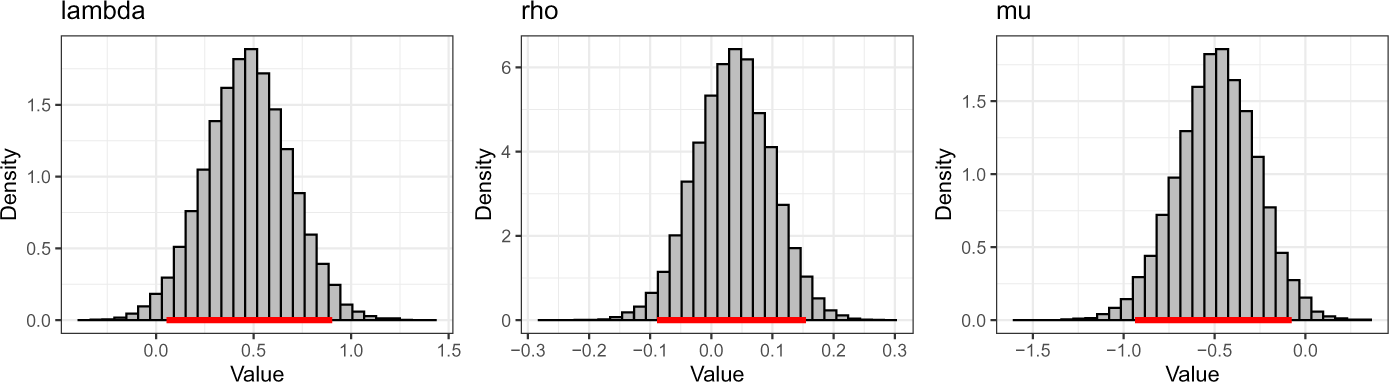
95% HDI of the group mean differences for the loss aversion, risk aversion and inverse temperature parameters of Model 1.

## G HDDMs - Parameter recovery

**Table 10:** Parameter recovery for the atvz, atz and tz models. We report Pearsons’ *r* correlations.

| atvz model | a | t | v | z |
| --- | --- | --- | --- | --- |
| High Anxiety - Mixed-gamble | 0.50 | 0.85 | 0.55 | 0.24 |
| High Anxiety - Gain-only | 0.06 | 0.65 | 0.33 | 0.41 |
| Low Anxiety - Mixed-gamble | 0.27 | 0.89 | 0.51 | 0.13 |
| Low Anxiety - Gain-only | 0.21 | 0.85 | 0.50 | 0.34 |

| atz model | a | t | v | z |
| --- | --- | --- | --- | --- |
| High Anxiety - Mixed-gamble | 0.14 | 0.85 | - | 0.35 |
| High Anxiety - Gain-only | 0.41 | 0.81 | - | 0.28 |
| Low Anxiety - Mixed-gamble | 0.53 | 0.89 | - | 0.62 |
| Low Anxiety - Gain-only | 0.24 | 0.77 | - | 0.36 |

| tz model | a | t | v | z |
| --- | --- | --- | --- | --- |
| High Anxiety - Mixed-gamble | - | 0.82 | - | 0.54 |
| High Anxiety - Gain-only | - | 0.81 | - | 0.13 |
| Low Anxiety - Mixed-gamble | - | 0.91 | - | 0.39 |
| Low Anxiety - Gain-only | - | 0.83 | - | 0.25 |

**Figure 6:**
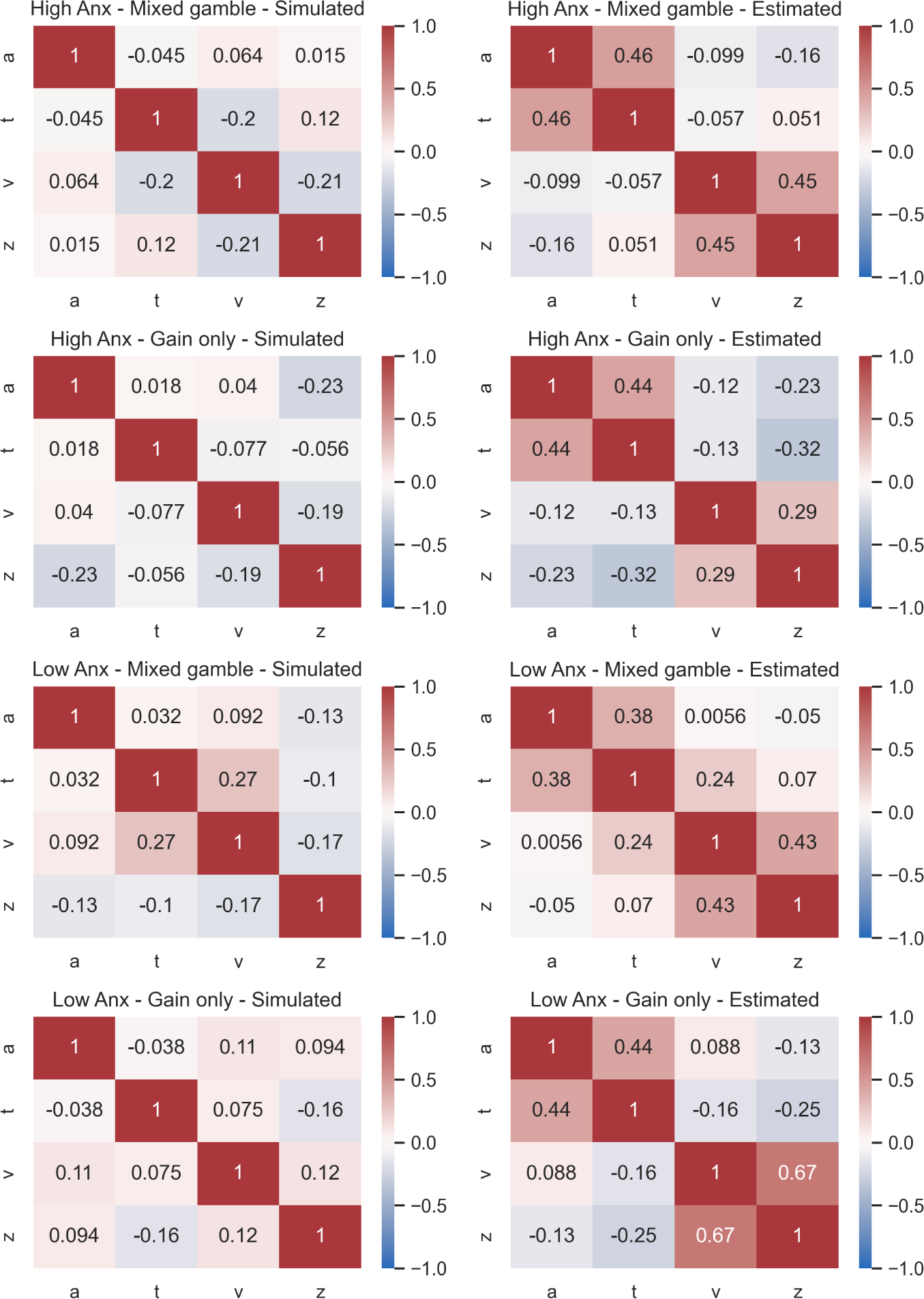
Correlations between parameters in the parameter recovery of the atvz model (best fitting model).

**Figure 7:**
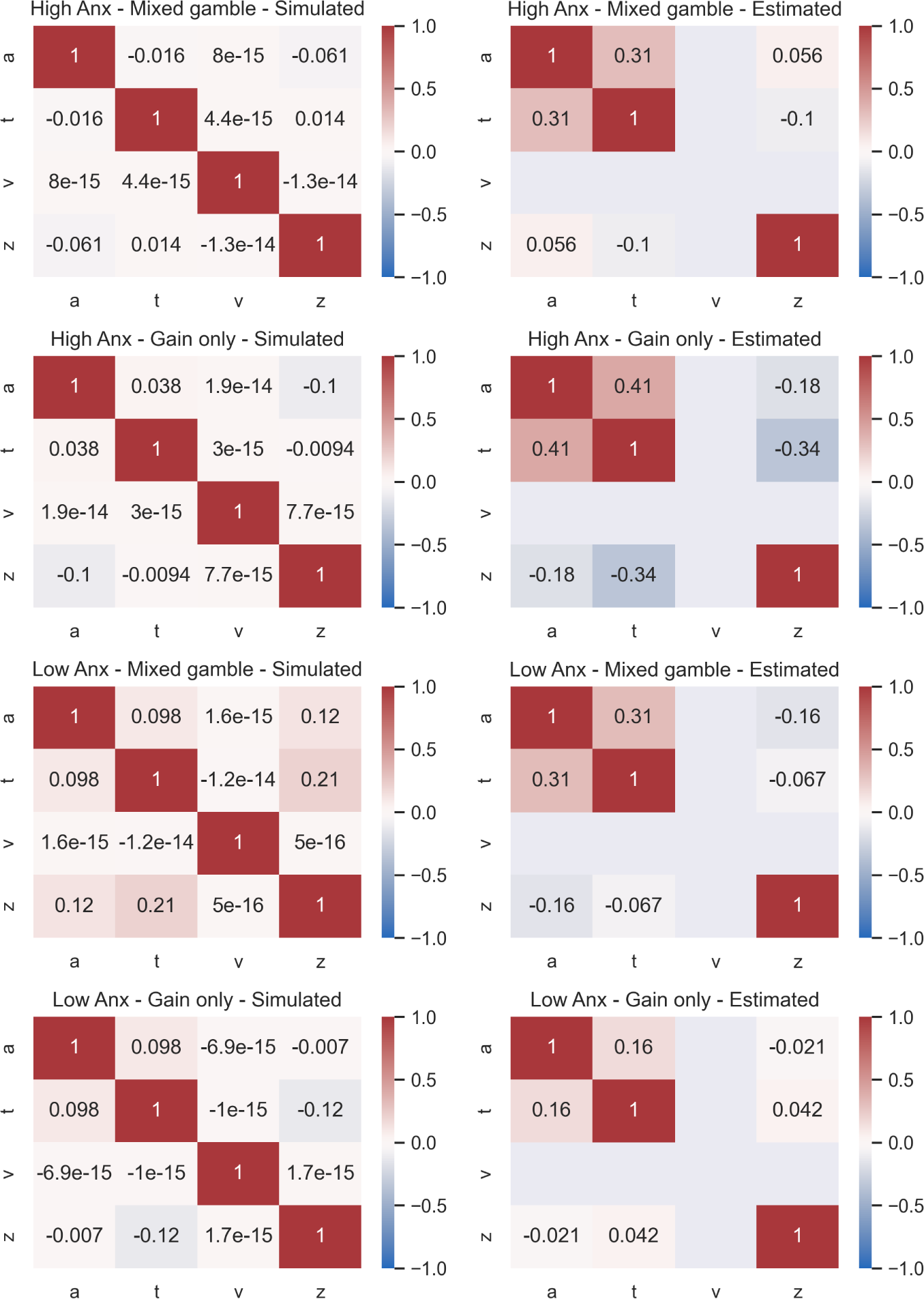
Correlations between parameters in the parameter recovery of the atz model (5th best fitting model).

**Figure 8:**
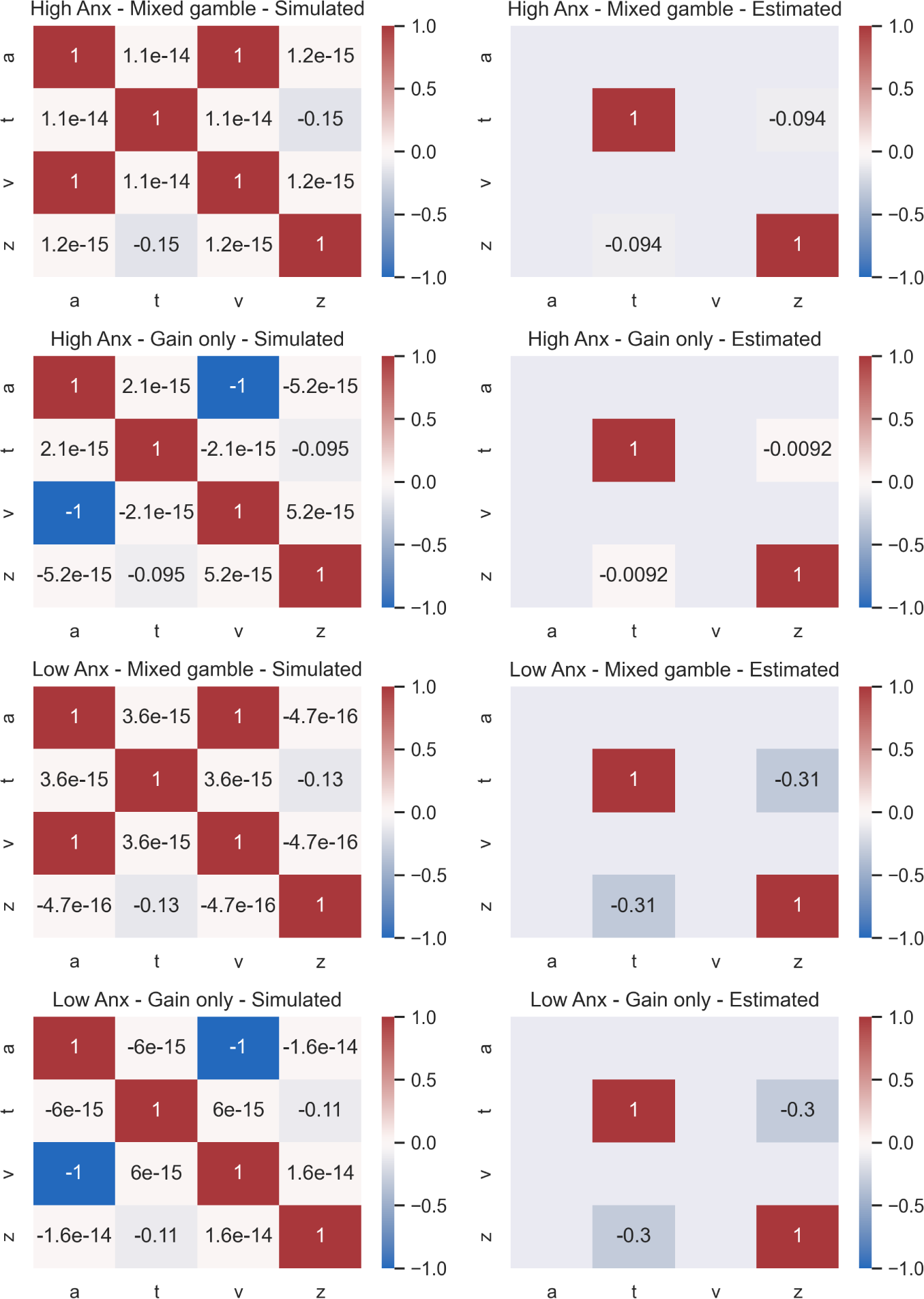
Correlations between parameters in the parameter recovery of the tz model (6th best fitting model).

## H HDDMs - Extra results

The model fitting results of the HDDMs are reported in table 11.

**Table 11:** Model fitting results based on Deviance Information Criterion (DIC), lower is better. a is boundary separation, t is non-decision time, v is drift-rate, z is starting point.

| Model | Low Anx DIC | High Anx DIC | Total DIC |
| --- | --- | --- | --- |
| **none** | 17071.3 | 15590.0 | 32661.3 |
| **a** | 16958.1 | 15468.9 | 32427.0 |
| **t** | 16182.5 | 15094.7 | 31277.2 |
| **v** | 16273.1 | 15183.7 | 31456.8 |
| **z** | 16453.7 | 15327.6 | 31781.3 |
| **at** | 16122.5 | 15045.0 | 31167.4 |
| **av** | 16238.9 | 15069.6 | 31308.5 |
| **az** | 16439.5 | 15240.9 | 31680.4 |
| **tv** | 15492.1 | 14699.7 | 30191.7 |
| **tz** | 15830.0 | 14890.8 | 30720.8 |
| **vz** | 16120.3 | 15079.1 | 31199.4 |
| **atv** | 15400.0 | 14660.7 | 30060.7 |
| **atz** | 15738.3 | 14858.0 | 30596.2 |
| **avz** | 16105.2 | 15002.5 | 31107.8 |
| **tvz** | 15450.8 | 14660.6 | 30111.3 |
| **atvz** | 15338.0 | 14601.5 | **29939.6** |

The correlations of the estimated parameters of models atv, tvz and tv are:

- atv: a and v are correlated in mixed-gamble trials (*r*_115_ = 0.408, *p <* 0.001), t is correlated with v in gain-only trials (*r*_115_ = *−*0.202, *p* = 0.030).
- tvz: v and z are correlated in mixed-gamble trials (*r*_115_ = 0.317, *p <* 0.001), t and v are correlated in gain-only trials (*r*_115_ = *−*0.219, *p* = 0.018).
- tv: t and v are correlated in gain-only trials (*r*_115_ = *−*0.200, *p* = 0.031).

The posterior distributions of the estimated parameters for the atvz model are shown in fig. 9.

**Figure 9:**
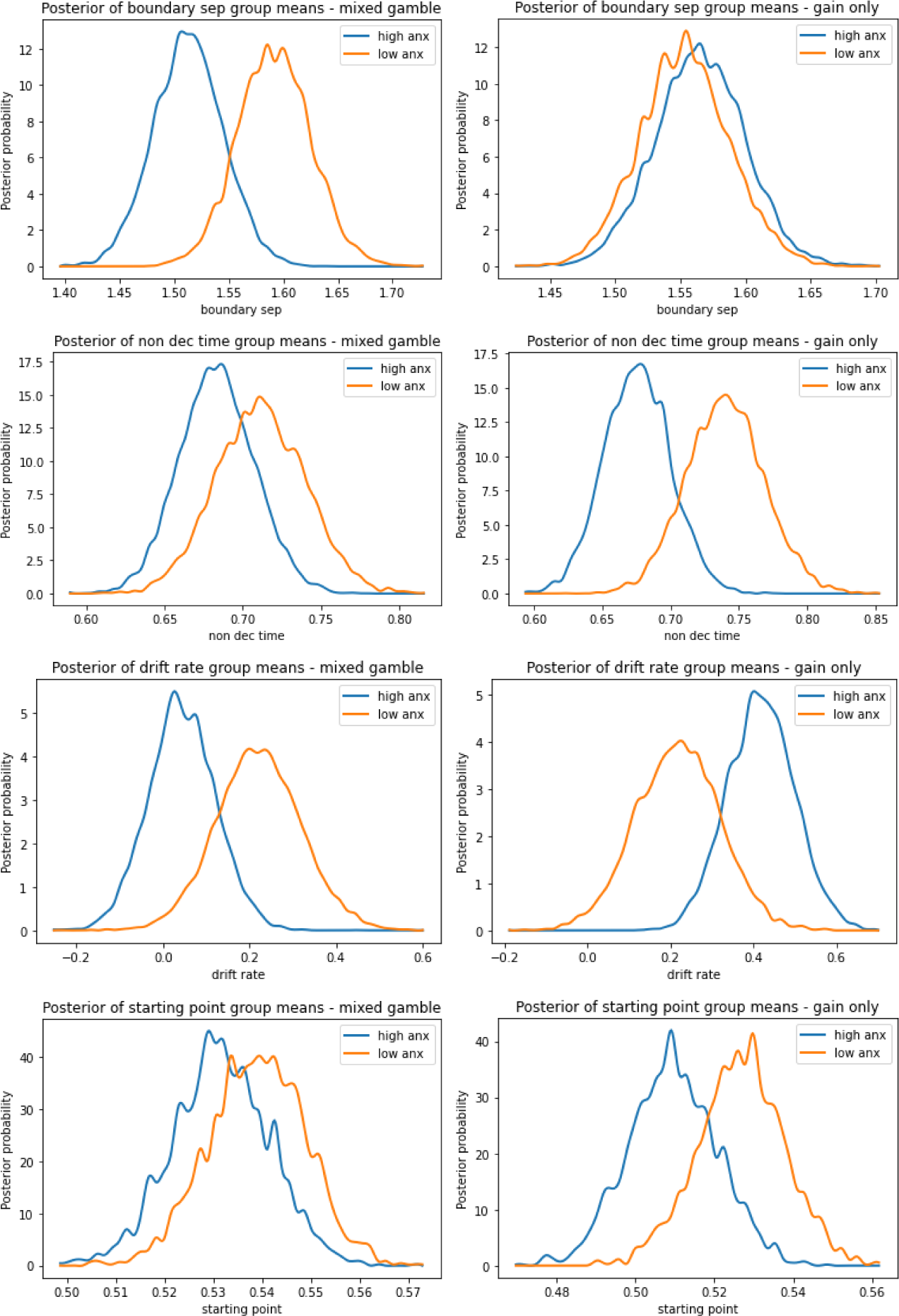
Posteriors distributions of the parameters fitted using the atvz model. Posteriors for the mixed-gamble trials are on the left.

